# Reference Gene Selection for Accurate RT-qPCR Normalization in Four Tissues and Whole-Body Samples of *Acheta domesticus*

**DOI:** 10.64898/2025.12.16.694577

**Authors:** Houda Ben-Miled, Nicolas Périard, Fanny Renois, Marie-Hélène Deschamps, François Meurens, Marie-Odile Benoit-Biancamano

## Abstract

House crickets (*Acheta domesticus*) are increasingly recognized as a sustainable protein source for food and feed systems. However, despite their growing relevance, molecular research on this species remains extremely limited, particularly concerning robust normalization strategies for gene expression analysis. This study is the first to identify and validate suitable reference genes for RT-qPCR analysis in *A. domesticus* across different tissues, an essential step for accurate quantification of host and pathogen target gene expression. Six candidate reference genes commonly used in insects (*AdoNEOPT, EF2, 18S rRNA, EF1α, Histone H3,* and *GAPDH*) were evaluated for expression stability in five tissue types (abdomen, legs, wings, head, and whole body). Gene stability was assessed using five computational tools: BestKeeper, geNorm, NormFinder, Delta Ct, and the integrated platform RefFinder. Additional validation was performed using the R statistical software. The results identified *EF1α*, *AdoNEOPT*, *EF2*, and *18S rRNA* as the most stable reference genes across all the selected tissues, while *GAPDH* and *His H3* showed high variability and were generally unsuitable except in the head, where *GAPDH* demonstrated stable expression. This study provides the first validated set of reference genes for *A. domesticus*, laying a foundation for accurate and reproducible gene expression studies. Moreover, our study will enable the development of new diagnostic tests based on qPCR and molecular signatures. These tests will be essential tools for health monitoring in insect farms, which remain exposed to emerging diseases.

## Introduction

Global pressures on food systems including the growing demand for protein, climate change, and biodiversity loss are accelerating the development of insect farming as a sustainable complement to conventional livestock production (Carthy et al., 2018). Crickets are particularly promising for large-scale production of protein due to their high feed conversion efficiency, relatively low land and water requirements, and their varied applications in human food, animal feed, and bioproducts (Takacs et al., 2023).

The house cricket (*Acheta domesticus*) is among the most widely farmed insect species and has also been established as a model organism for studying physiology, immunity, and environmental adaptation under intensive farming conditions. Having been reared for decades across Europe, North America, and Asia, it is now increasingly utilized by the agri-food industry as an ingredient in a variety of food and feed products (Osimani et al., 2018; Udomsil et al., 2019). As with other species subjected to intensive rearing, *A. domesticus* populations are exposed to numerous biotic and abiotic stressors, including pathogens, overcrowding, dietary shifts, and temperature fluctuations, factors that can adversely affect growth, reproduction, and survival (Vaga et al., 2021; Pilco-Romero et al., 2023). Gaining insights into the molecular mechanisms underpinning these stress responses is therefore crucial for optimizing health management and ensuring sustainable productivity in commercial cricket farming operations. Reverse transcriptase-quantitative polymerase chain reaction (RT-qPCR) remains the benchmark technique for quantifying transcript abundance, owing to its high sensitivity, specificity, and broad accessibility (Schmittgen and Livak, 2008). However, its accuracy is critically dependent on the use of stably expressed reference genes for normalization. These internal controls are essential for correcting variability in RNA quality and quantity, as well as in reverse transcription efficiency, across different tissues and experimental conditions (Kozera and Rapacz, 2013). However, numerous studies have demonstrated that the expression of commonly used reference genes is not inherently stable (Basu et al., 2019; Jeon et al., 2020; Lee et al., 2025). Therefore, choosing appropriate reference genes is crucial for obtaining reliable results of gene expression analyses.

Despite recent advances in genomics and biotechnology in *A. domesticus* (Oppert et al., 2020; Dossey et al., 2023; Homchan et al., 2024), rigorous evaluations of candidate reference genes, differentiated by tissue type, is still missing. Many studies still rely on traditional reference genes, not even available in *A. domesticus*, without prior validation (see discussion) an approach that can introduce significant biases and lead to misinterpretation of expression dynamics, particularly when comparing different tissues or under various stress conditions relevant to farming and health monitoring.

The transcription levels of commonly used reference genes such as 18S ribosomal RNA (*18S rRNA*), elongation factor 1-alpha (*EF1α*), and glyceraldehyde-3-phosphate dehydrogenase (*GAPDH*) (Jeon et al., 2020; Kim & Kim, 2025; Kozera & Rapacz, 2013; Zhao et al., 2022), have long been presumed to be uniformly expressed and unaffected by experimental conditions, given their essential roles in fundamental cellular processes (Kozera and Rapacz, 2013). As a result, they have traditionally been employed as single reference genes for normalization. However, multiple studies have demonstrated that their expression can vary and may be unstable under different experimental setups (Lin and Lai, 2010; Chapuis et al., 2011; Lü et al., 2018; Jeon et al., 2020; Khan et al., 2022).

In this study, we systematically evaluated the expression stability of several candidate reference genes in *A. domesticus,* including *18S rRNA, EF1α, GAPDH, AdoNEOPT,* histone H3 *(His H3*) and elongation factor 2 (*EF2*) across different tissues (head, wings, legs, abdomen) as well as in whole-body samples. We incorporated multiple complementary algorithms geNorm, NormFinder, BestKeeper, RefFinder, and the Delta cycle threshold (ΔCt) method in addition to analyzing the cycle quantification (Cq) or Ct value distribution, to rank the stability of the candidate genes. We also used pairwise variation analysis to determine the optimal number of reference genes required for robust normalization based on the tissue analyzed. The recommendations from this study provide a practical and validated framework for RT-qPCR normalization in *A. domesticus*, enabling more reliable host and pathogen gene expression studies in support of health monitoring, experimental physiology, and the overall optimization of house cricket farming systems.

## Materials and methods

### Insects

Live specimens of house crickets (*A. domesticus*) were obtained from a healthy commercial colony (MagaZoo, Montreal, Quebec, Canada), exhibiting no evidence of morbidity, mortality, or clinical signs such as melanization, wing deformities, or external body lesions, and transported to the laboratory in clean, ventilated boxes. They were then immediately frozen at −80°C until RNA extraction.

### RNA extraction and cDNA synthesis

Adult house crickets (*A. domesticus*) were used for the analysis of reference gene expression. Four tissue types: abdomen, head, legs, and wings were dissected separately, in addition to whole-body samples (Figure 1). Antennae were removed from the head samples. Each sample type was prepared using ten biological replicates. Whole body and tissue (abdomen, head, legs, wings) images of *A. domesticus* were captured using a Moticam S3 camera (Motic, Vancouver, Canada) and a ring light (LED 60T-B, Bausch & Lomb, Rochester, USA), both mounted on a dissection stereomicroscope (SMZ 161 Series, Leica Microsystems, Buffalo Grove, Chicago, USA).

**Figure 1:**
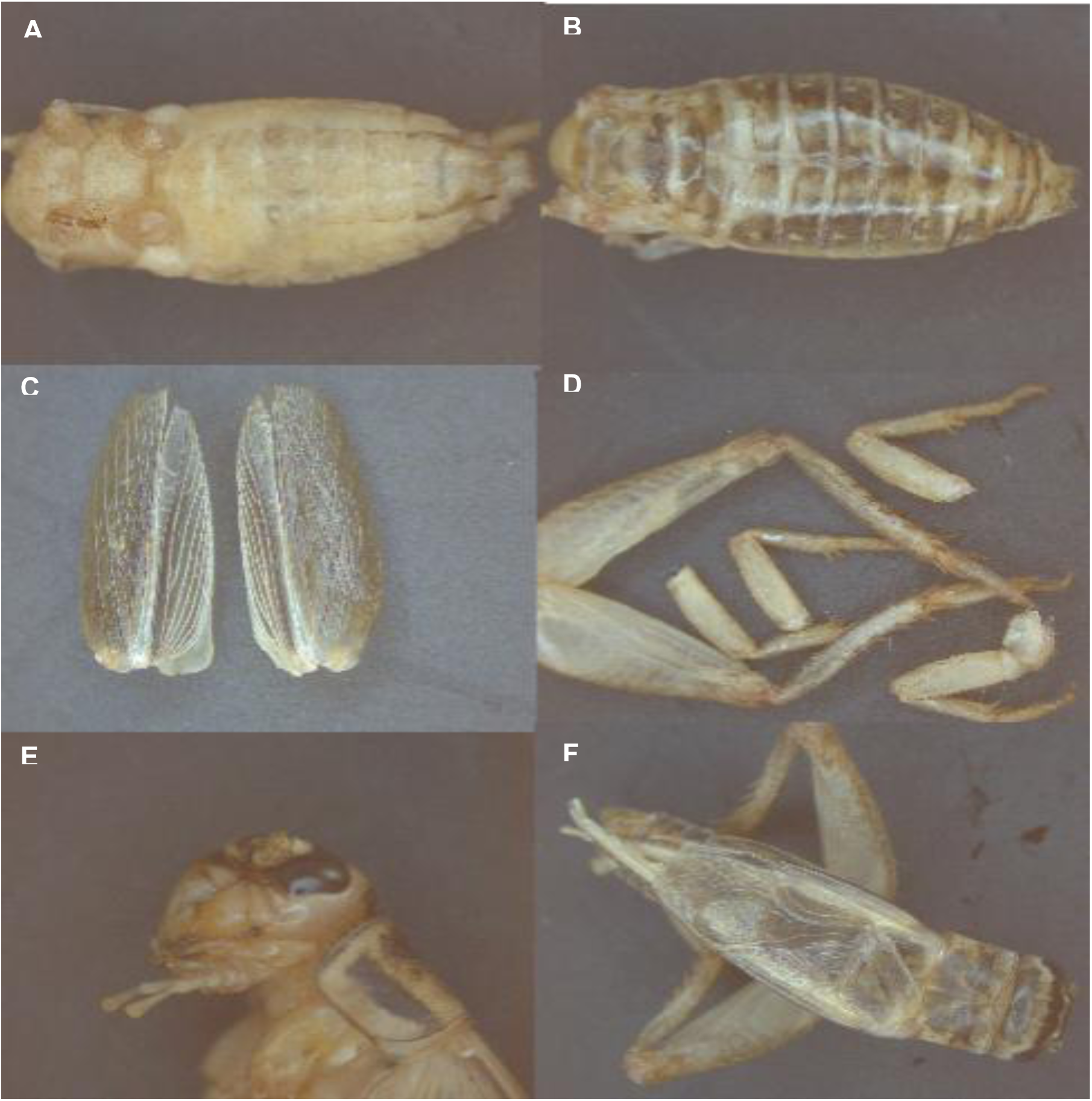
Macroscopic views of adult *Acheta domesticus*. (A, B) Ventral perspectives (A) abdominal view, (B) dorsal view; (C) wings view; (D) legs view; (E) head view; and (F) whole-body view. Images acquired at a magnification between 7.5× and 10× (objective: 0.75–1×; ocular: 10×).

RNA extraction was performed using the RNeasy Plus Mini Kit (Qiagen, Stockach, Germany), with protocol adaptations depending on the tissue type. Tissues were individually placed in sterile Eppendorf (Thermo Fisher Scientific, Toronto, Canada) tubes containing 800 µL of RLT buffer and 15 µL of proteinase K (Qiagen, Stockach, Germany) for whole-body and abdominal samples (Figure 1F, A and B), and 600 µL of RLT buffer and 10 µL of proteinase K for head, wing, and leg samples (Figure 1E, C and D). Samples were ground using a sterile plastic pestle (Thermo Fisher Scientific, Toronto, Canada), ensuring complete homogenization. Following homogenization, the tubes were incubated for 6 hours at 60 °C. The resulting liquid homogenate was transferred to a new sterile tube for RNA purification, which was performed using the RNeasy Mini Kit (Qiagen, Stockach, Germany) with on-column DNase treatment, according to the manufacturer’s instructions. RNA was eluted in 30 µL of RNase-free water and quantified using a NanoDrop™ One/OneC spectrophotometer (Thermo Fisher Scientific, San Diego, CA, USA). Only samples with an A260/280 ratio between 1.8 and 2.1 and an A260/230 ratio greater than 2 were selected for further analysis (Bustin et al., 2025). RNA was stored at −80 °C until further use.

For each sample, 500 ng of total RNA were used for cDNA synthesis using the iScript™ Advanced cDNA Synthesis Kit for RT-qPCR (Bio-Rad, Hercules, CA, USA). Reverse transcription reactions were performed under the following conditions: 5 min at 25 °C, 20 min at 46 °C, and 1 min at 95 °C. The resulting cDNA was diluted 1:10 in RNase-free water and stored at −20 °C for subsequent qPCR analysis.

### Selection of candidate genes and primer design

Candidate gene selection was based on annotated transcript sequences of *A. domesticus* available in the NCBI transcriptomic databases (https://www.ncbi.nlm.nih.gov/) and the full genome (GenBank: JAHLJT010004567.1), as well as on reference genes commonly used in other insect species as reported in previous studies (Lü et al., 2018). BLAST and tBLASTn searches were conducted to identify expressed sequence tags (ESTs) in *Acheta domesticus*. For the BLAST analyses, we used gene sequences from phylogenetically related insects, particularly other cricket species such as *Acrididae*, *Locusta migratoria* (Q. Yang et al., 2014; Foquet and Song, 2020). These sequences allowed us to target homologous regions in *Acheta*, thereby facilitating the identification of candidate sequences. The tBLASTn searches, on the other hand, were performed using known protein sequences from other model insects (such as grasshoppers (L. Zhao et al., 2021)), enabling us to query available EST databases for *Acheta domesticus* as it was carried out in other species too (Bruel et al., 2010; L. Zhao et al., 2021). This approach allowed us to identify ESTs corresponding to genes of interest, with the aim of designing primers for subsequent qPCR analyses.

Nine primer pairs were designed from the CDS sequences of the target genes using Clone Manager 9 software (Scientific & Educational Software, Cary, NC, USA), and two additional pairs were selected from the scientific literature (Table 1). Primer design was carried out rigorously, following standard RT-qPCR guidelines (Bustin et al., 2025): primer lengths between 18 and 22 nucleotides, annealing temperatures between 56 °C and 60 °C, and amplicon sizes ranging from 100 to 250 bp. Primers were synthesized by Integrated DNA Technologies (IDT, Ontario, Canada). The specificity of each primer pair was verified by qPCR using cDNA as a template. Amplification efficiency (E) for each primer pair was calculated using the formula: E = (10^^[–1/slope]^ – 1) × 100, where the slope was obtained from a standard curve generated using a 5-fold serial dilution of cDNA templates. In total, six primer pairs were selected for subsequent experiments. The six selected reference genes were: *GAPDH*, *EF2*, *EF1α*, *AdoNEOPT*, *His H3*, and *18S rRNA*. These genes were selected because of their essential roles in distinct cellular processes and the reported stability of their expression in insects, particularly in *Acheta domesticus* and other *Orthoptera* species (Zhu et al., 2014; Lü et al., 2018; Lu et al., 2022; Takacs et al., 2023). The sequences, amplicon sizes, and amplification efficiencies of the primers used are presented in Table 1.

**Table 1:**
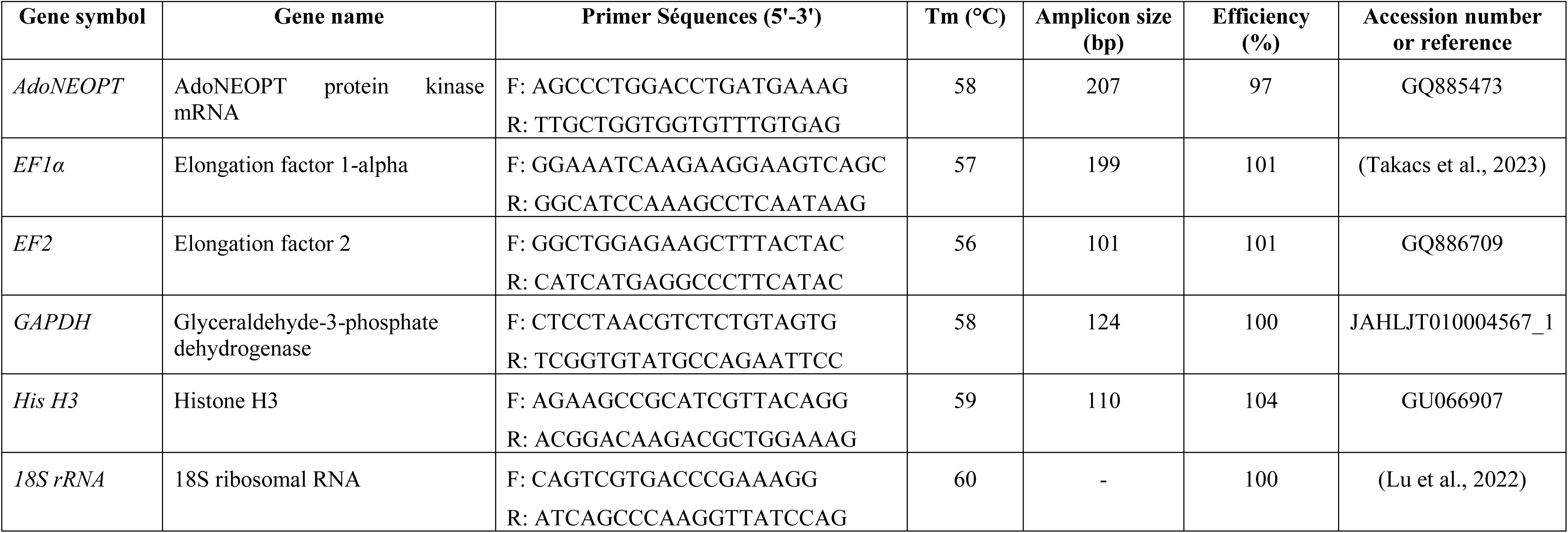
Features of candidate reference genes in *Acheta domesticus*.

### Quantitative Real-Time PCR

The CFX Opus 96 real-time PCR detection system (Bio-Rad, Hercules, USA) in combination with the SsoAdvanced Universal SYBR Green Supermix chemistry. qPCR analysis was carried out using ten biological replicates and two technical replicates per sample, in a final reaction volume of 10 μL containing 2 μL of synthesized cDNA (diluted 1:10), 5 μL of SsoAdvanced Universal SYBR Green Supermix (2X, Bio-Rad, Hercules, CA, USA), 0.2 μL of each primer (forward and reverse; IDT, Ontario, Canada), and nuclease-free water to adjust the final volume to 10 μL. A no-template control (NTC) was included to check for contamination of the reaction mix. qPCR reactions were performed under the following thermal cycling conditions: an initial activation step at 98 °C for 30 seconds, followed by 40 cycles of denaturation at 98 °C for 5 seconds and annealing/extension at 55–60 °C for 30 seconds. At the end of the amplification, a melt curve analysis (from 65 °C to 95 °C in 0.5 °C increments) (Figure S1) was performed to verify the specificity of the amplicons. The Cq for each well was determined using the CFX Maestro software (version 2.3, 2001) with a single Cq (Ct) threshold determination mode.

### Analyzing Reference Genes and Handling Data

To determine the expression stability of six candidate reference genes, we used RefFinder, an online tool that integrates four algorithms: geNorm (version 3.5) (Vandesompele et al., 2002), NormFinder (version 0.953) (Andersen et al., 2004), BestKeeper (version 1) (Pfaffl et al., 2004), and the delta cycle threshold (ΔCt) method (Silver et al., 2006). In addition, RefFinder (Xie et al., 2012) (https://www.ciidirsinaloa.com.mx/RefFinder-master/) integrates and consolidates the results from these different methods to provide a comprehensive ranking of candidate genes and identify the most stable reference genes. The geNorm software calculates the gene expression stability value (M) for each candidate gene and generates a ranking, where genes with the lowest M values are considered the most stable (Vandesompele et al., 2002). NormFinder software identifies the most appropriate reference genes for normalization. It calculates an expression stability value for each candidate gene, with lower values indicating greater stability (Andersen et al., 2004). The BestKeeper algorithm identifies stable reference genes based on minimal variation in Cq values. It calculates parameters such as the geometric mean, arithmetic mean, standard deviation (SD), and coefficient of variation (CV), and performs correlation analyses (R²) between each candidate gene and the others. Candidate genes that show strong correlations are then combined to assess *p*-values. According to BestKeeper, candidate genes with standard deviation (SD) values lower than 1 and a relatively low coefficient of variation are considered to be more stable reference genes. A standard deviation greater than 1 is regarded as indicative of unstable expression (Chechi et al., 2012; Jeon et al., 2020; Y. Liu et al., 2023).

In addition, R software (version 4.5; (Team, 2016)) was used to calculate pairwise variation (Vₙ/Vₙ₊₁), which measures the difference between two sequential normalization factors to determine the optimal number of reference genes required for accurate normalization. A Vₙ/Vₙ₊₁ ratio below 0.15 indicates that adding another reference gene would not significantly improve normalization (Vandesompele et al., 2002). The delta Ct method assesses gene expression stability by comparing the mean differences in Ct values between pairs of genes. By using each candidate gene as a reference, the standard deviations of the delta Ct values of the remaining genes are calculated to identify the most stable gene. The distribution of Cq values for the candidate genes across different body parts, as well as in whole-body cricket samples, was analyzed using the arithmetic mean (AM), standard deviation (SD), and coefficient of variation (CV), calculated as CV = SD/AM.

### Validation of reference genes

Expression profiles of the most stable reference gene, normalized either by a single reference gene or by multiple reference genes, were determined from cycle threshold (Cq) values using the 2^−ΔΔCt^ method (Livak and Schmittgen, 2001). Normality of the data was assessed using Anderson-Darling, D’Agostino & Pearson, and Shapiro-Wilk tests. A one-way analysis of variance (ANOVA), followed by Tukey’s post hoc test (p ≤ 0.05), was performed to compare expression means. Statistical analyses were conducted using GraphPad Prism 8 software (GraphPad, San Diego, CA, USA).

## Results

### Specificity and Amplification Efficiency of Candidate Reference Genes

Before performing RT-qPCR, the specificity and efficiency of the primers designed for the candidate reference genes were carefully evaluated. A total of six genes (*GAPDH, EF2, EF1α, AdoNEOPT, His H3,* and *18S rRNA*) were selected for analysis. All primer pairs generated a single peak in the melting curves (Figure S1), with no indication of nonspecific products or primer-dimer formation. No signal was detected in the negative controls, confirming the specificity of the reactions. The amplification efficiencies of the primers ranged from 100% to 107%, with regression coefficients (R²) greater than 0.996, thus meeting the quality criteria required for qPCR analyses. Detailed information on the genes, primer sequences, amplicon sizes, annealing temperatures (Tm), and efficiencies (E) is presented in Table 1.

### Distribution of Cq Values of Reference Genes

In this study, the Cq values of each candidate reference gene were measured in cDNA samples samples of the house cricket (*A. domesticus*) collected from four tissue types (head, wings, legs, abdomen), as well as the whole body. To highlight differences in transcription levels among the six candidate reference genes, the stability of Cq values plays a crucial role in selecting reference genes; the expression level of a gene depends on its Cq value; the lower the Cq value, the higher the expression level, and vice versa.

Furthermore, the Cq values of the candidate reference genes varied depending on the tissue type. Therefore, it is necessary to analyze the range of Cq values and calculate the coefficient of variation (CV) for each gene across all samples. Figure 2 shows that the Cq values of the six reference genes ranged from 8.11 (*18S rRNA*) to 30.58 (*GAPDH*), with 80% of the values falling between 20 and 29 cycles.

**Figure 2:**
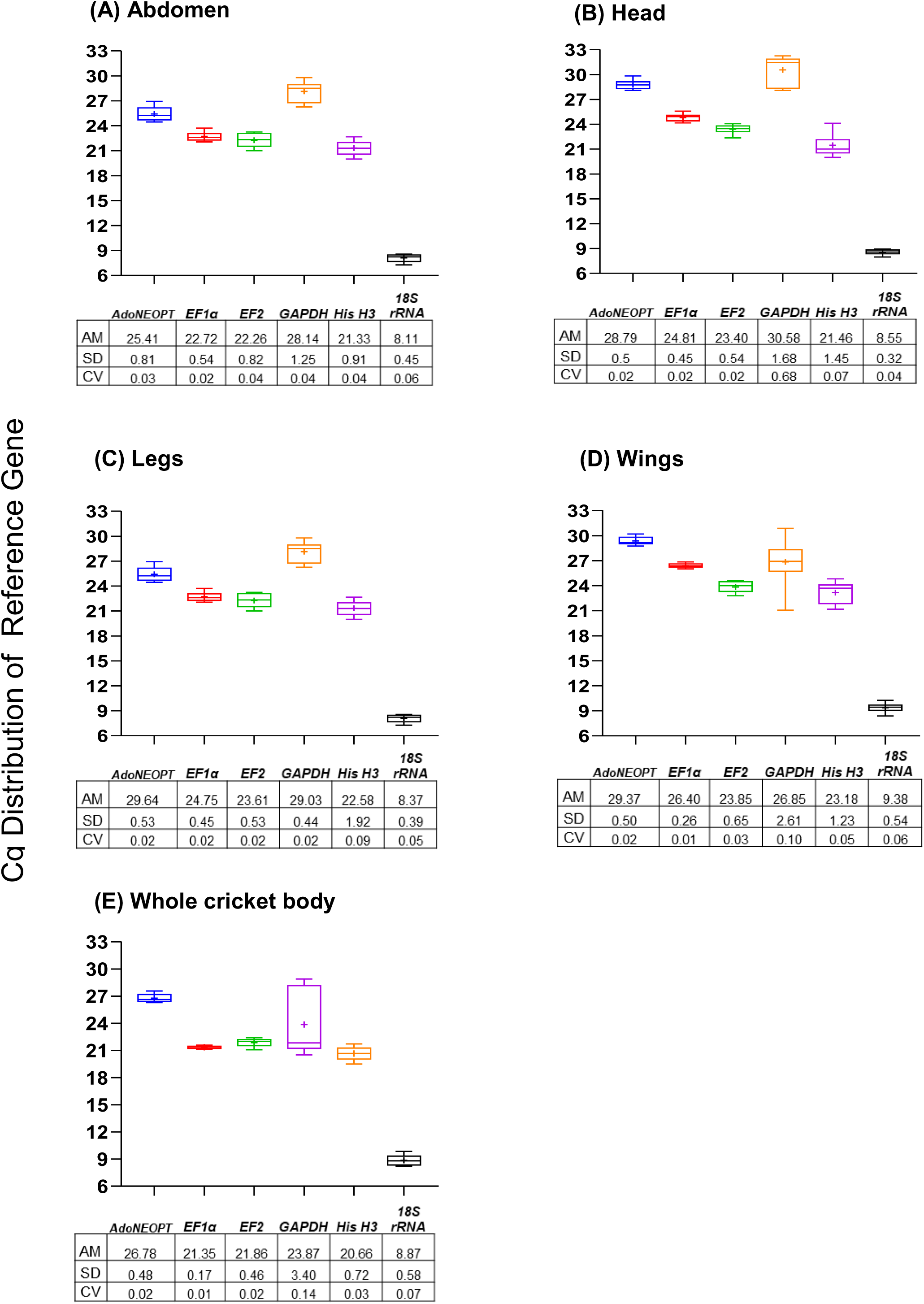
Box plot comparisons of Cq values for the six candidate reference genes in criket (*Acheta domesticus*) samples. Samples were prepared from four body compartments of *Acheta domesticus* (A–D), whole cricket body (E), The horizontal lines inside the boxes indicate the 25th, 50th (median), and 75th percentiles. The cross symbol inside the main box represents the mean. Error bars denote the minimum and maximum values. AR: Arithmetic Mean, SD: Standard Deviation, CV: Coefficient of variation.

Expression stability was also assessed by calculating the coefficient of variation (CV) of the Cq values, with CV < 1 being a criterion indicating low variance (Ospina & Marmolejo-Ramos, 2019). *EF1α* was also the most stable gene, with the lowest CV values (0.02) in the abdomen, wings, and whole body, whereas *GAPDH* showed the greatest variation, with a CV reaching 0.10 in the wings and whole body (Figure 2A, D and E).

In the head and legs, *EF1α, AdoNEOPT*, and *EF2* showed the lowest CVs (0.02), indicating very stable expression (Figure 2B and C). In contrast, *His H3* in the head was found to be less stable, with the highest CV of 0.09 (Figure 2B).

### Expression Stability of Candidate Reference Genes

To determine the stability and rank the candidate reference genes, we used geNorm, NormFinder, BestKeeper, delta-Ct, and R.

#### geNorm analysis

The geNorm analysis was used to rank the reference genes by calculating the gene expression stability value (M) based on the average pairwise variation of expression ratios. The most stable reference gene has the lowest M value, while the least stable gene has the highest M value. As suggested by previous studies, an M value < 1 is considered an acceptable threshold to classify the tested genes as relatively stable (Hellemans et al., 2007; H. Kim & Kim, 2025; Liu et al., 2014).

By comparing M values across tissue types, *EF1α, AdoNEOPT, EF2*, and *18S rRNA* were identified as reliable reference genes, all showing M values < 1. The results showed that the *EF1α + AdoNEOPT* pair exhibited the highest stability in the abdomen and whole body, while *EF1α + 18S rRNA* was the most stable combination in the wings. The two most stable genes in the legs were *EF1α* and *GAPDH*, while the most stable pair in the head was *AdoNEOPT* and *EF2*. In contrast, *GAPDH* showed the least stable expression in the abdomen, wings, head, and whole body of the house cricket, with M values ≥ 1. Similarly, *His H3* displayed marked instability in the legs and head, with an M value of 1. Conversely, *GAPDH* showed very stable expression in the legs (M = 0.3), and *His H3* was also found to be stable (M < 1) in the abdomen, wings, and whole body (Figure 3 and Table 3).

**Figure 3:**
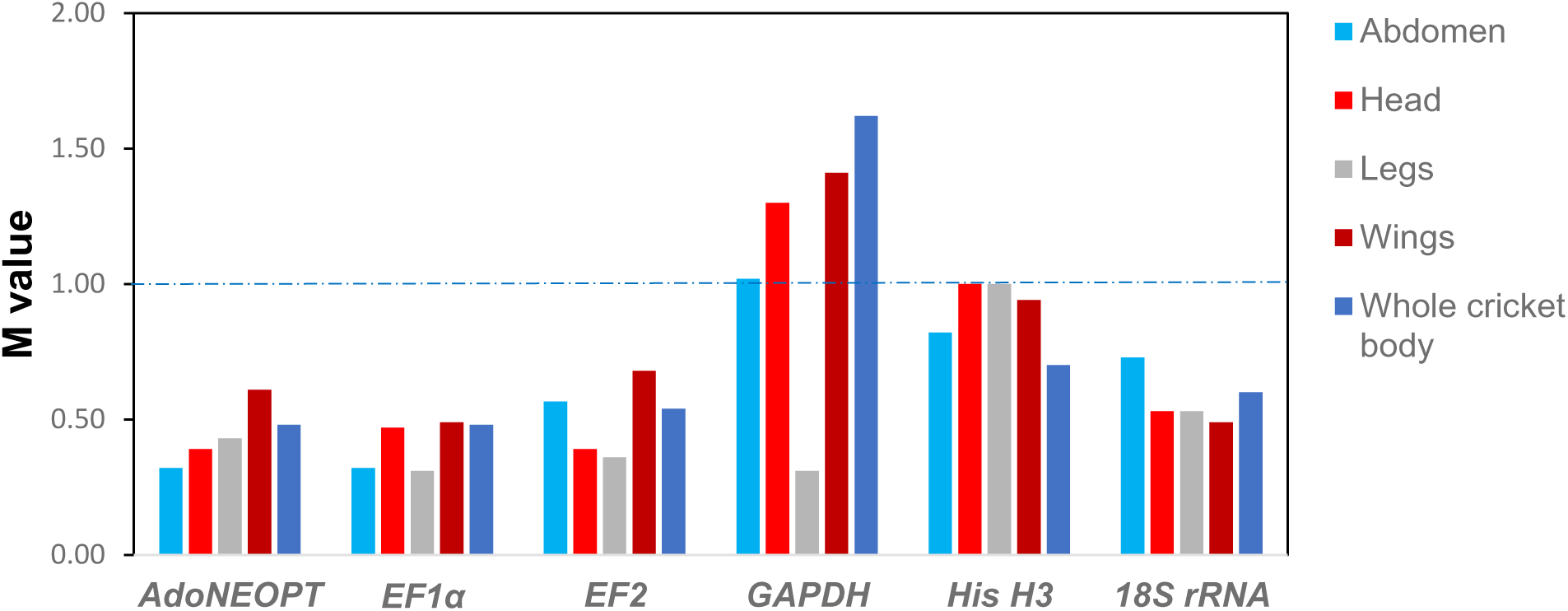
Average expression stability (M) values calculated by geNorm for six candidate reference genes in *Acheta domesticus* across four body compartments and the whole cricket body.

Thus, there was little variation in the ranking of reference genes across the four tissues studied and the whole body. *EF1α* showed the lowest M value, indicating the highest expression stability in all samples, except in the head. Overall, the analysis confirmed that all tested reference genes exhibited M values below the acceptable threshold, except for *GAPDH* and *His H3*, which showed variable stability depending on the tissue type.

#### NormFinder Analysis

The NormFinder software evaluates the reliability of reference genes based on both intra- and inter-group variations. The most stable genes are those with the lowest stability values. However, no specific threshold value is recommended by this program. Figure and Table present the stability values of the reference genes for each tissue type. The results showed that *EF1α* was the most reliable reference gene in the abdomen, legs, and whole body, while *AdoNEOPT* was the most stable in the wings. Finally, *18S rRNA* was identified as the optimal gene for the head. According to the analysis performed with NormFinder, *GAPDH* was found to be the least stable gene across all samples, except in the legs (Figure 4).

**Figure 4:**
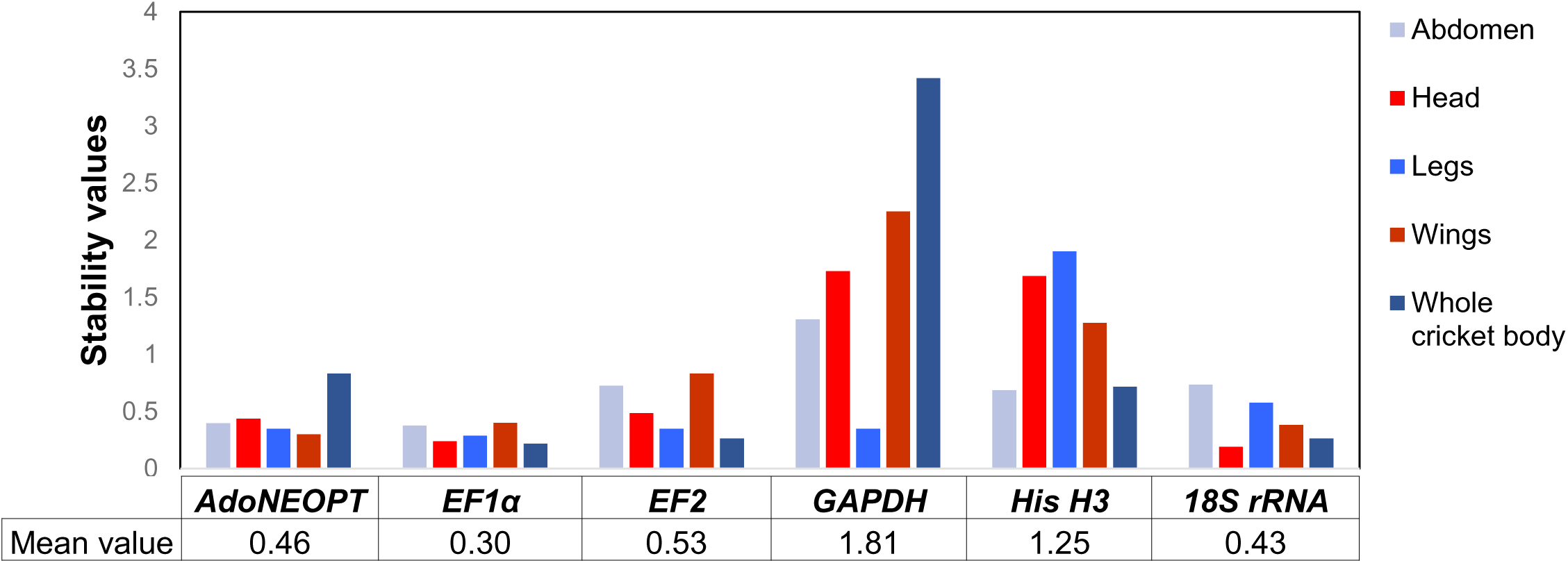
Expression stability values of the six candidate reference genes in *Acheta domesticus*, calculated using NormFinder. Average stability values were arithmetically calculated based on data from four body compartments and the whole body of the cricket.

#### BestKeeper Analysis

Gene expression variation was calculated for the six candidate reference genes based on their Cq values, and expressed as standard deviation (SD) and coefficient of variation (CV) using the BestKeeper software. Lower SD values (generally < 1) indicate more stable expression, allowing the corresponding genes to be considered the most stable. BestKeeper identified *18S rRNA* as the reference gene with the lowest overall variation in the abdomen, legs, and head among the six candidates, with an SD of 0.3. *EF1α* was found to be the most stable gene in the wings and whole body, with SD values of 0.14 and 0, respectively (Table 2).

**Table 2:** Gene expression stability values of the six candidate reference genes analyzed using BestKeeper.

| Rank | Abdomen |  |  |  |  |  | Head |  |  |  |  |  | Legs |  |  |  |  |  |
| --- | --- | --- | --- | --- | --- | --- | --- | --- | --- | --- | --- | --- | --- | --- | --- | --- | --- | --- |
|  | Gene | SD<sup>a</sup> | CV<sup>b</sup> | GM<sup>c</sup> | CD(R<sup>2</sup>)<sup>d</sup> | *p* value | Gene | SD<sup>a</sup> | CV<sup>b</sup> | GM<sup>c</sup> | CD(R<sup>2</sup>)<sup>d</sup> | *p* value | Gene | SD<sup>a</sup> | CV<sup>b</sup> | GM<sup>c</sup> | CD(R<sup>2</sup>)<sup>d</sup> | *p* value |
| 1 | *18S rRNA* | 0.33 | 4.1 | 7.96 | 0.006 | 0.81 | *18S rRNA* | 0.3 | 3.61 | 8.42 | 0.309 | 0.095 | *18S rRNA* | 0.36 | 4.56 | 7.88 | 0.41 | 0.046 |
| 2 | *EF1α* | 0.49 | 2.17 | 22.47 | 0.233 | 0.15 | *EF2* | 0.39 | 1.7 | 23.39 | 0.081 | 0.425 | *EF2* | 0.42 | 1.8 | 23.3 | 0.312 | 0.093 |
| 3 | *AdoNEOPT* | 0.51 | 2.02 | 25.3 | 0.338 | 0.07 | *EF1α* | 0.4 | 1.63 | 24.68 | 0.077 | 0.437 | *AdoNEOPT* | 0.42 | 1.43 | 29.3 | 0.175 | 0.228 |
| 4 | *EF2* | 0.64 | 2.91 | 22.19 | 0.488 | 0.03 | *AdoNEOPT* | 0.48 | 1.69 | 28.71 | 0.203 | 0.19 | *GAPDH* | 0.42 | 1.48 | 28.3 | 0.068 | 0.467 |
| 5 | *His H3* | 0.66 | 3.11 | 21.24 | 0.273 | 0.12 | *His H3* | 1.17 | 5.51 | 21.22 | 0.313 | 0.093 | *EF1α* | 0.48 | 1.97 | 24.4 | 0.253 | 0.138 |
| 6 | *GAPDH* | 0.97 | 3.47 | 27.89 | 0.043 | 0.56 | *GAPDH* | 1.4 | 4.6 | 30.41 | 0.156 | 0.258 | *His H3* | 1.6 | 7.21 | 22.13 | 0.407 | 0.047 |
| Rank | Wings |  |  |  |  |  | Whole cricket body |  |  |  |  |  |  |  |  |  |  |  |
|  | Gene | SD<sup>a</sup> | CV<sup>b</sup> | GM<sup>c</sup> | CD(R<sup>2</sup>)<sup>d</sup> | *p* value | Gene | SD<sup>a</sup> | CV<sup>b</sup> | GM<sup>c</sup> | CD(R<sup>2</sup>)<sup>d</sup> | *p* value |  |  |  |  |  |  |
| 1 | *EF1α* | 0.14 | 0.55 | 26.1 | 0.372 | 0.057 | *EF1α* | 0 | 0 | 21 | 0 | 0.364 |  |  |  |  |  |  |
| 2 | *18S rRNA* | 0.45 | 4.84 | 9.28 | 0.504 | 0.019 | *AdoNEOPT* | 0.42 | 1.6 | 26.3 | 0 | 0.091 |  |  |  |  |  |  |
| 3 | *AdoNEOPT* | 0.48 | 1.66 | 29.2 | 0.608 | 0.007 | *18S rRNA* | 0.48 | 5.71 | 8.39 | 0.315 | 0.18 |  |  |  |  |  |  |
| 4 | *EF2* | 0.59 | 2.49 | 23.63 | 0.048 | 0.537 | *EF2* | 0.5 | 2.33 | 21.49 | 0.212 | 0.291 |  |  |  |  |  |  |
| 5 | *His H3* | 1.08 | 4.67 | 23.13 | 0.16 | 0.243 | *His H3* | 0.64 | 3.17 | 20.19 | 0.137 | 0.001 |  |  |  |  |  |  |
| 6 | *GAPDH* | 1.67 | 6.27 | 26.64 | 0.756 | 0.001 | *GAPDH* | 3.08 | 13.16 | 23.18 | 0.77 | 0 |  |  |  |  |  |  |

<sup>a</sup>SD indicates the standard deviation of Cq values.<sup>b</sup>CV refers to the coefficient of variation value.<sup>c</sup>GM represents the geometric mean of Cq values.<sup>d</sup>CD (R<sup>2</sup>) indicates the coefficient of determination value.

Although gene stability varied across tissues, all six genes analyzed had SD < 1.0 in the abdomen, suggesting that each of them could potentially serve as a reference gene for normalizing target gene expression in this anatomical region of the house cricket (Table 2). For the other tissues, all genes also had SD < 1.0, except for *GAPDH*, which showed SD > 1 in the wings, head, and whole body, and *His H3*, which had SD > 1 in the head, wings, and legs. The results from the BestKeeper analysis showed some differences compared to those obtained with geNorm and NormFinder (Table 3).

**Table 3:**
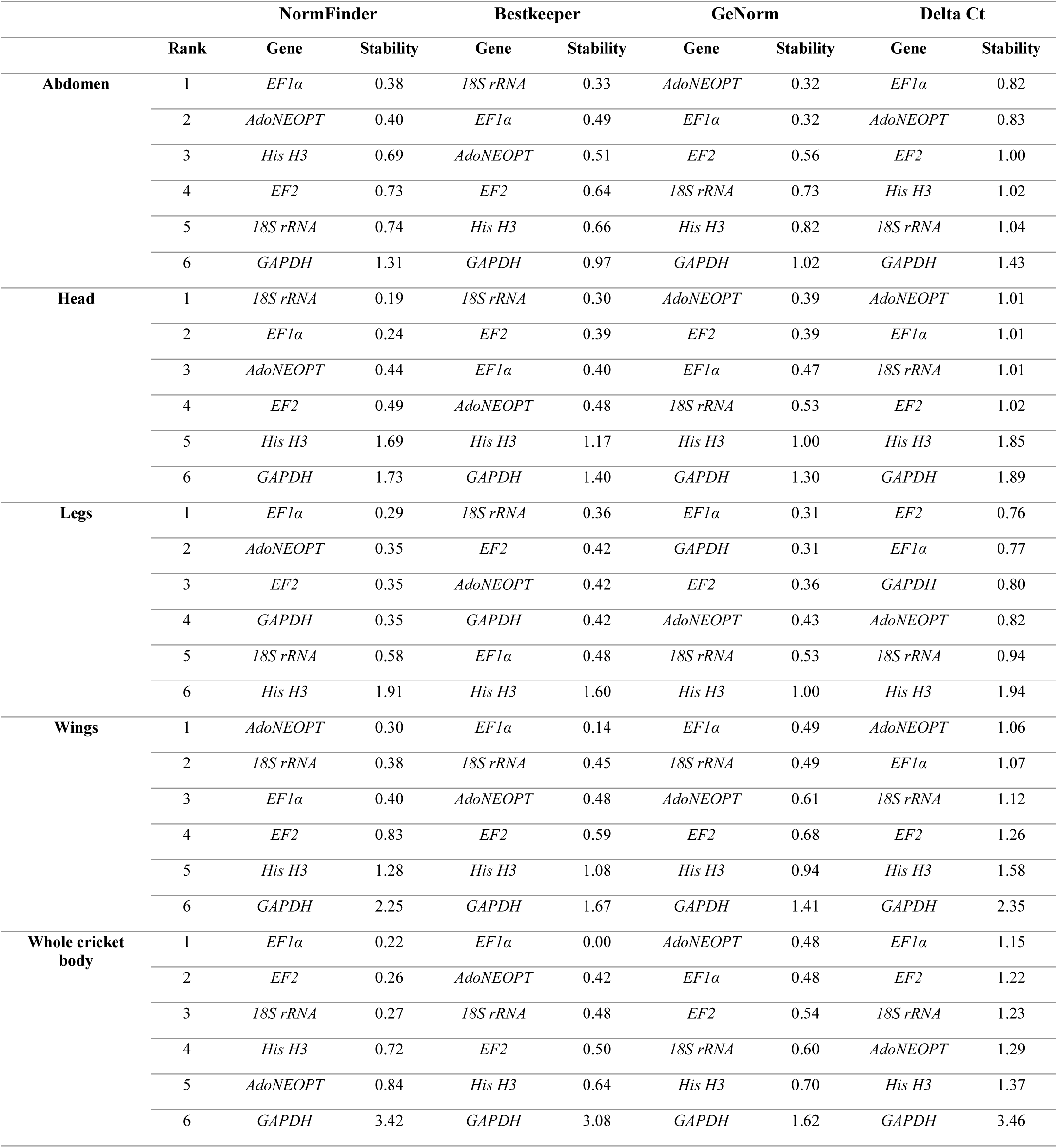
Stability ranking of six candidate reference genes across five body parts using four different algorithms.

#### Delta-Ct Analysis

In this approach, gene expression stability is assessed by calculating the mean and standard deviation (SD) for each gene. The comparative ΔCt analysis revealed that *GAPDH* showed the least stable expression across the different tissues, except in the legs, where it appeared to be stable. *EF1α* proved to be the best gene for normalization in the abdomen, head, and whole body. The most reliable gene in the wings was *AdoNEOPT*, while the most stable gene in the legs was *EF2* (Table 3).

#### Pairwise Variation Analysis Using R software

To determine the optimal number of reference genes required for accurate normalization, pairwise variation (V_n_/V_n+1_) was calculated using the R software. In general, a threshold value of 0.15 is used to identify the optimal number of reference genes.

Our results showed that, for the head and legs, the V2/3 value was below 0.15, indicating that two reference genes are sufficient for gene expression normalization in these tissues (Figure 5). In contrast, for the wings, the V3/4 value was below 0.15, suggesting that three reference genes are necessary for reliable normalization. For the abdomen, the V5/6 value remained above 0.15, indicating that more than six reference genes may be needed to ensure optimal normalization in this tissue.

**Figure 5:**
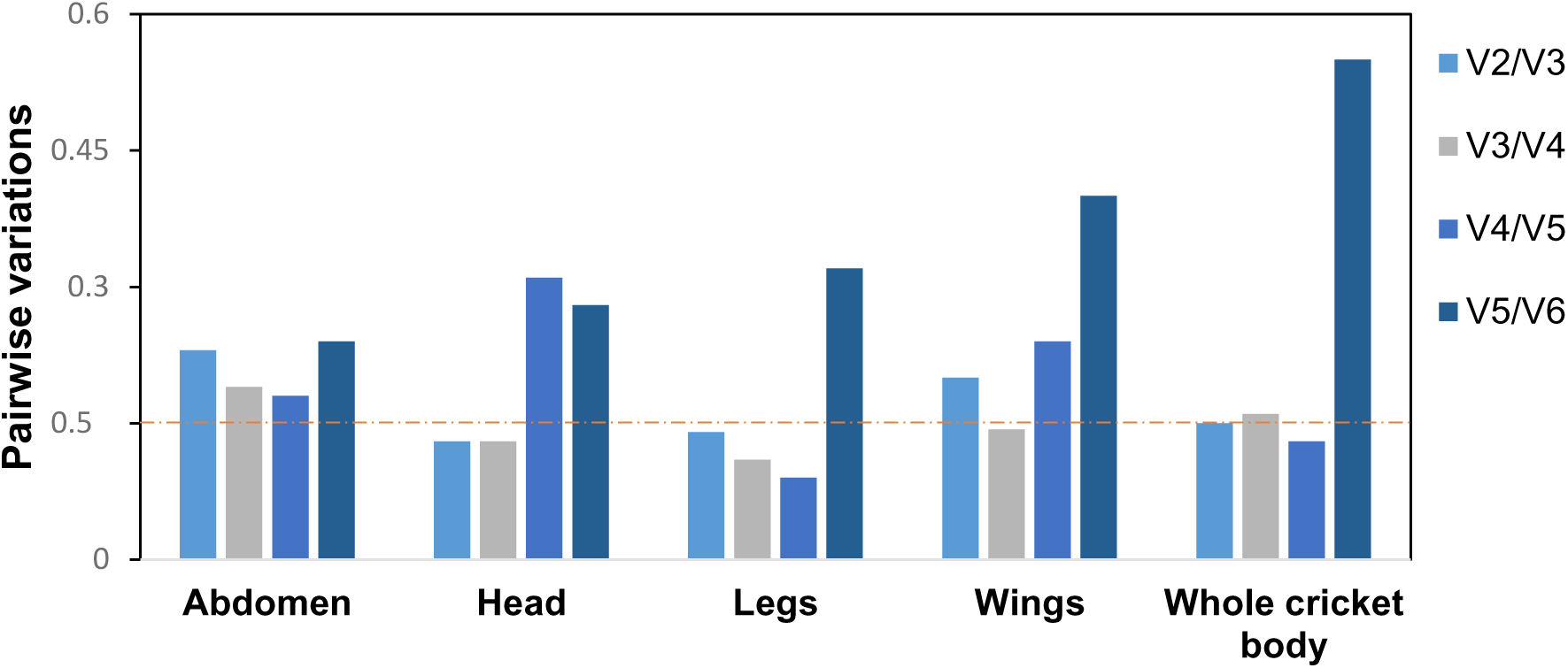
Pairwise variation analysis, performed using R software, was employed to determine the optimal number of reference genes required for target gene normalization. Pairwise variation values (Vn/Vn+1) were calculated from data obtained from four body compartments of *Acheta domesticus* and the whole cricket body. The dotted lines represent the pairwise variation threshold set at 0.15, which serves as the cutoff value to define the optimal number of reference genes.

Finally, for the whole body, the V2/3 value was exactly 0.15, while V4/5 was 0.13. This suggests that two genes might be sufficient, although this is right at the threshold. To be cautious, the use of four reference genes is therefore recommended to ensure reliable normalization in this case (Figure 5).

### Overall Ranking of Reference Gene Expression Stability According to RefFinder

Reference genes were ranked in the house cricket across different tissues using the online tool RefFinder, which integrates the results from three algorithms (NormFinder, BestKeeper, and geNorm). The overall expression stability of each gene was determined based on the geometric mean of its rank across all three algorithms, with stability increasing as the geometric mean decreases.

The results show that the reference gene *EF1α* was the most reliable in the abdomen, legs, wings, and whole body, while *AdoNEOPT* was the most stable gene in the head (Figure 6). Additionally, *AdoNEOPT* was ranked second in terms of stability in the abdomen, wings, and whole body, whereas *18S rRNA* was identified as the third most stable gene in the abdomen, wings, and whole body, and the second most stable in the head. In contrast, in the legs, *EF2* appeared as the second most stable gene (Figure 6A-E). In most tissues, *GAPDH* proved to be the least reliable reference gene.

**Figure 6:**
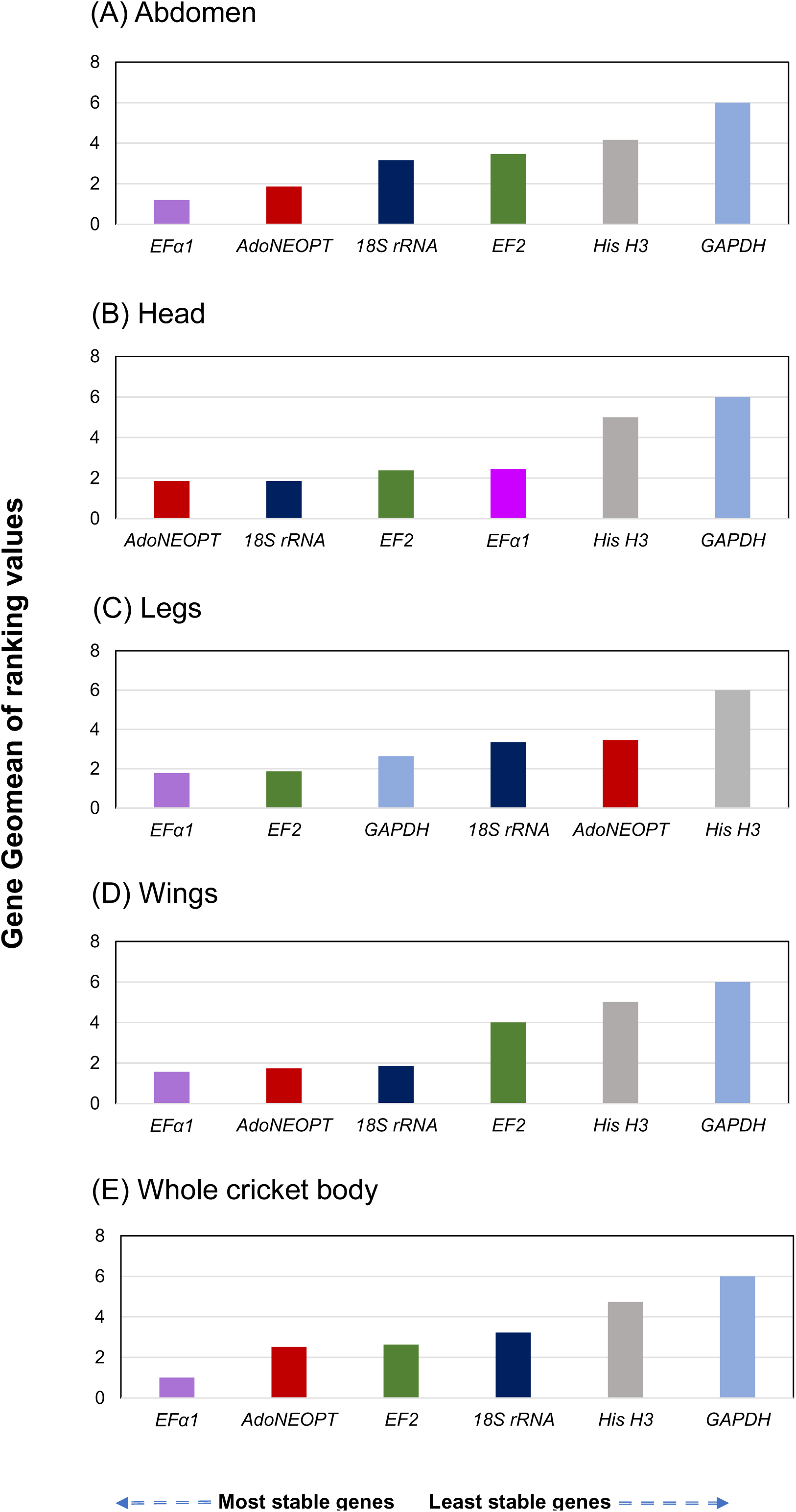
Expression stability of six candidate reference genes across different samples. The geometric method implemented in RefFinder was used to calculate the overall stability ranking of the reference genes. A lower Geomean ranking value indicates more stable expression. The four body compartments of *Acheta domesticus* (A–D) and the whole cricket body (E) were analyzed simultaneously.

### Validation of Selected Reference Genes

Based on the results obtained from the various analysis software (see above), EF1α was identified as the most stable reference gene across different body regions. It was therefore selected for the normalization of candidate gene expression. and its expression levels were compared when normalized using different candidate reference genes (*AdoNEOPT, EF2, GAPDH, His H3, 18S rRNA*) across various tissues as well as the whole body (Figure 7A-D). The normality of the data was assessed using several statistical tests, including Anderson-Darling, D’Agostino & Pearson, and Shapiro-Wilk. The results show that the data follow a normal distribution (*p* > 0.05), which justifies the use of parametric methods. This comparison was based on a rigorous statistical analysis using ANOVA, followed by Tukey’s post-hoc test. In the head, the expression of *EF1α* varied significantly depending on the reference gene used. *His H3* and *18S rRNA* yielded extreme and unreliable results, reflecting an overestimation and underestimation of expression, respectively. In contrast, *EF2* and *AdoNEOPT* provided normalization results more consistent with those obtained using *EF1α* itself, although statistically significant differences remained. The results also showed that variation in gene expression levels across different tissues was significant, regardless of the reference gene used, whether considered stable or not (P < 0.001 in all cases). However, *AdoNEOPT* and *EF2* produced more consistent and biologically plausible expression levels, indicating better normalization performance.

**Figure 7:**
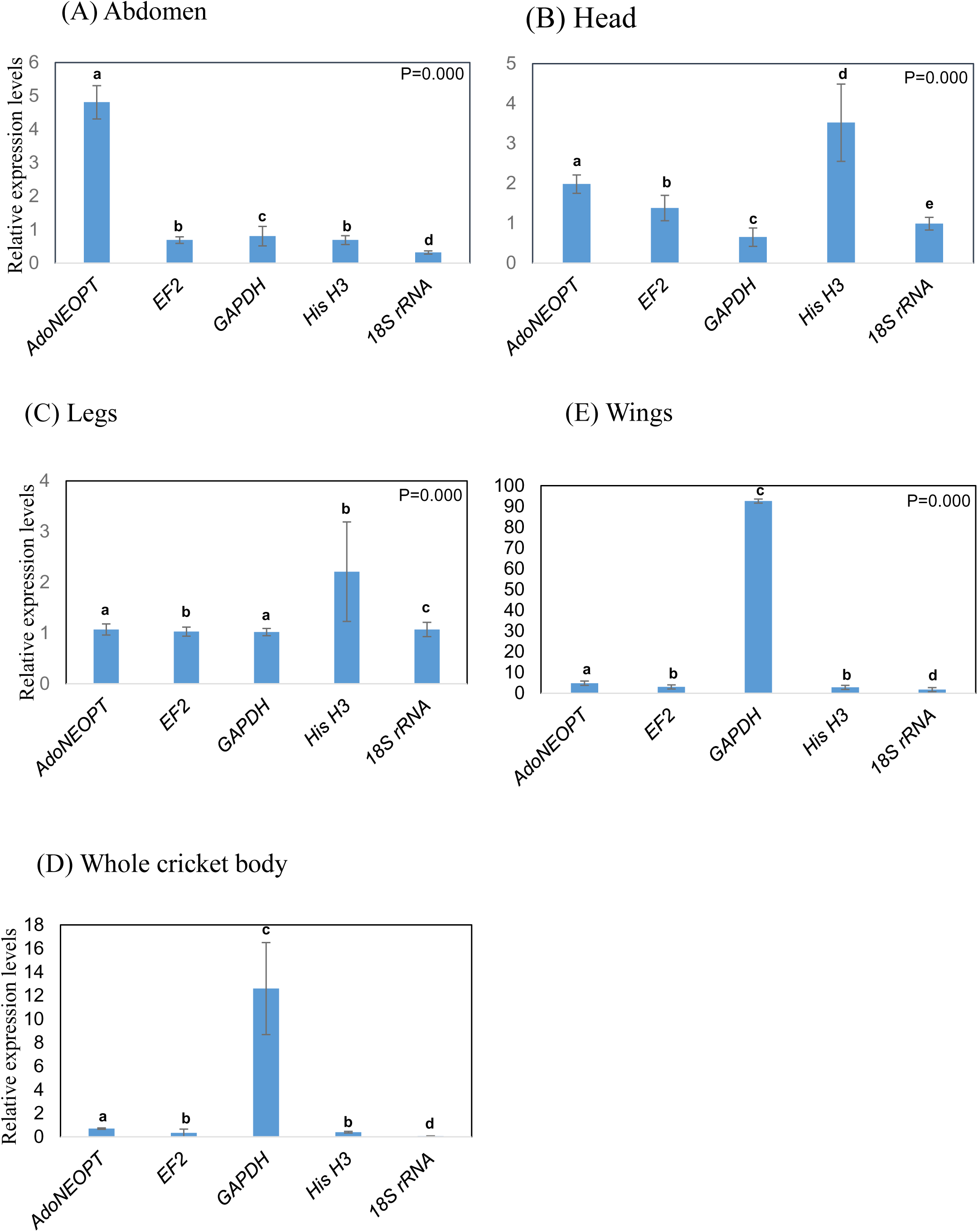
Comparison of expression levels of target genes in *Acheta domesticus* tissues normalised using the reference gene *EF1α*. Samples were prepared from for body part tissue (abdomen-A, head-B, legs-C, wings-E) and whole cricket body (D). Expression levels were statistically analysed using one-way ANOVA followed by Tukey’s multiple comparison post-hoc test, and different letters indicate significantly different values between genes within each tissue (*p*<0.05). Data are presented as mean values ± SE.

These observations highlight that the choice of reference gene greatly influences the estimation of target gene expression, and that prior validation of reference gene stability is essential for each specific tissue to ensure reliable and reproducible results.

## Discussion

RT-qPCR is a widely used technique in molecular biology for quantifying the expression of target genes across various tissues and experimental conditions (Lü et al., 2018). To ensure reliable and reproducible results, it is essential to identify stably expressed reference genes that enable accurate normalization of the data. However, no universally stable reference gene can be applied across all organisms and under all physiological or experimental conditions. Therefore, it is necessary to evaluate the expression stability of candidate genes (Khan et al., 2022). Despite a significant increase in publications on reference gene identification in insects (Omondi et al., 2015), this area remains underexplored in some *Orthoptera* species, particularly *A. domesticus*, which is amongst the most widely reared species for edible insects production intended for both human and animal consumption (Udomsil et al., 2019; Cohen, 2020).

To date, no study has been conducted to identify reliable reference genes in this species. This represents the first study aimed at evaluating and validating candidate reference genes in *A. domesticus*. Therefore, rigorous primer design is essential to avoid errors related to sequence polymorphisms at hybridization sites between species (Boyle et al., 2009). Indeed, even when primers are designed from genetically conserved regions, sequence differences can lead to variations in amplification efficiency, compromising the reliability of the results (Lee et al., 2025). In this context, there is no doubt that the selection of appropriate reference genes for *A. domesticus* is crucial to support research on this insect, whose nutritional and economic importance is rapidly growing.

The primers targeting the reference genes *EF1α, GAPDH, EF2, 18S rRNA, His H3*, and *AdoNEOPT* demonstrated good amplification efficiency. We then evaluated the expression stability of six candidate genes in *A. domesticus* across different body parts (abdomen, wings, head, legs) as well as in whole-body samples. To avoid the limitations associated with relying on a single analysis tool (Kim & Kim, 2025), we analyzed Cq values using four commonly applied methods (geNorm, NormFinder, BestKeeper, and ΔCt). These were complemented by the integrative platform RefFinder and additional statistical processing in R. As each method relies on distinct statistical principles, the ranking of candidate gene stability varied accordingly, a pattern frequently reported in previous studies (J. Wang et al., 2018; Moon et al., 2018; Jeon et al., 2020; Köhsler et al., 2020).

Considering the recommended thresholds for each tool (M < 1 for geNorm, SD < 1 for BestKeeper ((Hellemans et al., 2007; Y. Liu et al., 2023)), five out of the six candidate genes can be considered suitable as reference genes for gene expression analysis in the abdomen, legs, and whole body. In contrast, only four genes meet these criteria for the wings and head of *A. domesticus*. Although most studies have not established clear thresholds for stability in NormFinder and ΔCt analyses (Ponton et al., 2011; Reim et al., 2013; Moon et al., 2018), our findings are further supported by the Cq value distribution. The six candidate genes exhibited low expression variability, with coefficients of variation (CV) below 1, which is generally interpreted as an indicator of stable expression, in agreement with the observations reported by Ospina and collaborators (Ospina and Marmolejo-Ramos, 2019).

The genes identified as the most stable by the four software tools showed overall consistency, although minor differences in rankings were observed particularly in the wing samples. geNorm and BestKeeper produced an identical ranking of the four most stable genes: *EF1α, 18S rRNA, AdoNEOPT*, and *EF2*. NormFinder, however, ranked them as follows: *AdoNEOPT, 18S rRNA, EF1α*, and *EF2*. The ΔCt method yielded a similar ranking to NormFinder, with only a slight swap between the second and third positions. These variations are expected and can be naturally explained by the methodological differences among the algorithms used (Hou et al., 2023). Moreover, this variability is supported by previous studies conducted on honeybees. For instance, Moon and colaborators reported differences in the ranking of reference genes between nurse and forager bees (Moon et al., 2018), while Jeon and collaborators also observed divergent rankings across different tissues of *Apis mellifera* depending on the algorithms applied (Jeon et al., 2020). These findings reinforce the robustness of our results and highlight the importance of validating reference genes under diverse biological contexts.

The analysis of abdomen samples also revealed five stable candidate genes. geNorm and ΔCt produced identical rankings for the top three genes: *EF1α, AdoNEOPT*, and *EF2*. The fourth and fifth positions were occupied by *18S rRNA* and *His H3* in geNorm, whereas ΔCt reversed this order. NormFinder confirmed the same top two genes (*EF1α* and *AdoNEOPT*), while BestKeeper identified *18S rRNA* as the most stable, followed by *EF1α* and *AdoNEOPT* (Table 3). However, a clear convergence was observed in the whole-body samples, where *EF1α* was consistently ranked first by all four tools, underscoring its robustness as a reference gene in this tissue. This result is consistent with the findings of Kim and Kim (H. Kim and Kim, 2025), who also reported differences in the ranking of reference genes between the head and abdomen of *Apis mellifera*. In their study, the gene Ras-related protein Rab 1A (RAB1a) was identified as the most stable in the head according to *geNorm*, *NormFinder*, and *RefFinder*, whereas ADP ribosylation factor 1 (ARF1) was determined to be the most stable in the abdomen based on *geNorm* and *NormFinder*. These findings support our observations regarding the tissue-specific variability in gene expression stability (Kim & Kim, 2025). Furthermore, Jeon and collaborators, who evaluated the stability of several reference genes across three different tissues of *Apis mellifera*, reported similar trends to those observed in the present study. The use of different algorithms in their analysis also led to variations in gene rankings depending on the tissue type, confirming that such variability is methodologically expected and emphasizing the importance of experimental validation of reference genes under each specific biological condition (Jeon et al., 2020).

These discrepancies in gene stability rankings across different tissues are common when using multiple algorithms, due to their distinct calculation criteria. BestKeeper and geNorm, for example, rely on pairwise comparison approaches and assume that the genes being analyzed are not co-regulated (Øvergård et al., 2010). In contrast, NormFinder conducts a subgroup-based analysis, taking into account both intra-group and inter-group variation in expression levels when calculating gene stability values (Liu et al., 2014). The ΔCt method follows a similar approach to that described in the geNorm report by Vandesompele, in which “gene pairs” are directly compared (Silver et al., 2006).

Consistent with reports in *Locusta migratoria* (Yang et al., 2014), *Apis cerana* and *Apis mellifera* (Shao et al., 2024), *Aphis gossypii* (Ma et al., 2016), and other insect species (Li et al., 2013; Lee et al., 2025), our study found that it is challenging to identify universally suitable reference genes for qPCR analyses, as all selected reference genes showed notable variation in transcription levels across different tissues. Moreover, the top-ranked reference genes also varied depending on the analysis program used. To generate a final ranking, we used RefFinder, an integrated evaluation tool that calculates stability scores based on the geometric mean of rankings from multiple algorithms, thereby helping to overcome the limitations of relying on a single method (Vasu et al., 2024). A similar strategy has been applied in studies on various species, including *Leptocybe invasa* (Liu et al., 2023) and *Sclerodermus guani* (Zhao et al., 2024). According to RefFinder, *EF1α* was identified as the most stable reference gene in the abdomen, legs, wings, and whole-body samples, while *AdoNEOPT* was selected as the most suitable gene in the head.

The *EF1α* gene, which plays a key role in protein synthesis, functions as an essential translation elongation factor by facilitating the incorporation of amino acids into the polypeptide chain during mRNA translation (Saito and Ito, 2013). In our study, this gene ranked first in expression stability for samples from the abdomen, wings, legs, and whole body. However, it was placed fourth in the head, suggesting relatively lower stability in this specific tissue. These findings are consistent with those of Yang and collaborators (2014) in *Locusta migratoria*, where EF1α was identified as one of the most stable genes across various tissues and organs. Similarly, Chapuis and collaborators (2011) confirmed that *EF1α* is the most stable gene in the insect *Chortoicetes terminifera*. More recently, Zhao and collaborators also demonstrated that this gene is an optimal reference gene across different tissues in *Sclerodermus guani* (Zhao et al., 2024). In contrast, Freitas and collaborators, identified *EF1α* as one of the least stable genes in three stingless bee species tissues, based on several analysis tools (Freitas et al., 2019). These discrepancies between studies may be attributed to species-specific differences and variations in experimental conditions.

*AdoNEOPT* was used as a novel reference gene, identified from sequencing data (NCBI, 2016). The sequence corresponds to a fragment of messenger RNA (mRNA) that partially encodes a protein in *Acheta domesticus* (NCBI, 2016). *AdoNEOPT* encodes a regulatory kinase, an enzyme that transfers phosphate groups and is often involved in cellular signalling pathways (Regier et al., 2010). In this study, *AdoNEOPT* was identified as the most stable reference gene in the head, and the second most stable in the abdomen, wings, and whole body of the house cricket.

*EF2* (Elongation Factor 2) plays a crucial role in the translation process by catalyzing the binding of aminoacyl-tRNA to the ribosome’s acceptor site, a process that depends on *GTP* hydrolysis (Kaul et al., 2011). In this study, *EF2* was identified as the second most stable candidate reference gene in the legs of *Acheta domesticus*, and as the third most stable gene in the whole body and head, while ranking fourth in terms of stability in the wings and abdomen. Other studies on insect reference genes have also supported *EF2* as a suitable reference gene in *Orthoptera* (Zhu et al., 2014).

The 18S ribosomal RNA (*18S rRNA*), a component of ribosomal RNA, showed high expression levels across all samples, with very low Cq values (ranging from 7.26 to 10.26), indicating a high abundance of transcripts, this result is consistent with the observations of Zhao and collaborators in *Sclerodermus guani*, where similar Cq values (ranging from 6 to 11) were also reported (R. Zhao et al., 2024). It was therefore ranked among the most stable genes, occupying third place in the abdomen and wings, and second place in the head. Our study also revealed that *18S rRNA* was sufficiently stable according to all analysis tools, particularly with BestKeeper, comparable results were reported in *Aphis gossypii* (Ma et al., 2016) and in *Sclerodermus guani* (R. Zhao et al., 2024), reinforcing our observations. According to the literature, ribosomal protein genes such as *18S rRNA* are considered good reference genes due to their expression in all cells for the synthesis of new ribosomes (Shakeel et al., 2018). This gene has demonstrated high stability under various abiotic and biotic conditions in insects, suggesting that the ribosomal protein gene family could become a valuable source of reference genes (Dheda et al., 2004; Ma et al., 2016; Y. Liu et al., 2023). Several recent studies have validated ribosomal protein genes as reference genes across different insect tissues and have suggested that *18S rRNA* represents an ideal reference gene for normalizing RT-qPCR data (Bagnall & Kotze, 2010; Khan et al., 2022; Zhao et al., 2024). However, in contrast to our results, some studies have shown tissue-dependent variations in the expression of these genes. This issue of *18S rRNA* instability as a reference gene has also been observed in different insect species, such as *Bemisia tabaci* (Li et al., 2013; Liang et al., 2014) and *Danaus plexippus* (Pan et al., 2015).

The *GAPDH* gene, involved in glycolysis (Barber et al., 2005), is commonly used as an internal reference gene in humans, mice, pig, and various insect species. However, our results, obtained using all the analysis programs employed, showed that *GAPDH* is not a universally reliable reference gene across all tested tissues. It demonstrated sufficient expression stability only in the legs, where it ranked as the third most stable gene. Moreover, although *GAPDH* is frequently selected as one of the most stable genes under different experimental conditions in many insect species (Lü et al., 2018), several studies have highlighted its variable expression. For example, in species of the *Anopheles hyrcanus* group, its expression was found to be unstable across developmental stages (Lee et al., 2025). Additionally, Ma and collaborators study on *Aphis gossypii* also demonstrated that *GAPDH* is not a reliable reference gene under all experimental conditions (Ma et al., 2016). These observations underscore the importance of rigorously evaluating the stability of *GAPDH* in each specific context before using it as a reference gene in RT-qPCR analyses.

*His H3*, the reference gene encoding histone H3, a protein associated with chromatin structure (Elsaesser et al., 2010), was ranked among the two least suitable genes for normalization in the tested tissues according to our study. We found that *His H3* did not exhibit sufficient expression stability to be used as a reliable reference gene across different tissues. These results are consistent with other studies conducted on various insect species Yang et al., 2014), which also reported low stability of this gene in similar contexts. However, a study by Xu and collaborators on *Chilo suppressalis* showed contrary results, revealing *His H3* as the most stable gene among those evaluated across different tissues (Xu et al., 2017). This discrepancy highlights the importance of evaluating reference genes on a case-by-case basis, depending on the species, tissue, and experimental conditions.

This study highlights the importance of rigorous screening and selection of reference genes, a crucial step to ensure reliable and accurate normalization of gene expression data and the development or reliable diagnostic tests. As reported by several previous studies (Moon et al., 2018; Y. Kim et al., 2023), using a single reference gene can simplify technical procedures and reduce costs, but it may be insufficient under complex or variable biological conditions. Conversely, using too many reference genes can generate redundant data, complicate analyses and increase costs without a proportional gain in accuracy. Therefore, it is generally recommended to use two stable reference genes, as suggested by Majerowicz and colaborators (2011) and supported by other studies (Ponton et al., 2011; J. Wang et al., 2018; Y. Liu et al., 2023). According to the research by Vandesompele and collaborators (2002), which established a method to determine the optimal number of reference genes to use by calculating the pairwise variation factor V_n_/V_n+1_, a value below 0.15 is generally considered indicative of a sufficient number of genes for reliable normalization.

In this study, the number of reference genes required for RT-qPCR data normalization varied depending on the tissue type analyzed. For head and leg tissues, the V2/3 value was below 0.15, suggesting that two genes are sufficient for accurate normalization. In the wings, the V3/4 value was below 0.15, indicating that a set of three genes is necessary for adequate normalization. For whole body samples, the V2/3 value was exactly 0.15, while V4/5 was below this threshold, suggesting that using either two or four reference genes could be considered. This variability in the optimal number of reference genes according to the tissue studied is consistent with the findings of Kan and collaborators in *Paederus fuscipes* (Khan et al., 2022), who also demonstrated that the number of required genes depends on the type of tissue (abdomen, thorax, or head). These observations confirm that the selection of reference genes must be adapted to the biological and experimental context in order to ensure the reliability of gene expression analyses. However, due to the additional resources required to include four genes, using two genes is recommended, especially since the 0.15 threshold is not a strict rule. Some studies even accept slightly higher values (Wang et al., 2021; Zhao et al., 2022). Finally, in the abdomen, the V5/6 value exceeded the 0.15 threshold, indicating that at least six reference genes would be necessary for optimal normalization in this tissue.

These results demonstrate that combining multiple reference genes is essential to ensure reliable quantification of gene expression in *Acheta domesticus*, depending on the tissue studied. An appropriate selection, based on robust statistical analyses such as those provided by the various analysis programs, is therefore crucial to guarantee the validity of RT-qPCR results. In conclusion, our study paves the way for numerous further investigations focused on measuring the expression of *A. domesticus* transcripts or various pathogens affecting it. Rigorous normalization of qPCR results is crucial for developing and validating new qPCR-based diagnostic tools, as well as identifying molecular signatures characteristic of specific pathologies (e.g., various infections including emrging ones) or stress conditions.

## Supporting information

Melt curves

## Acknowledgments

This research was funded by the Ministère de l’Agriculture, des Pêcheries et de l’Alimentation du Québec (MAPAQ) through its Innov’Action Agroalimentaire program. François Meurens is supported by the Natural Science and Engineering Research Council of Canada (NSERC, grant RGPIN-2024-04212). CRIPA-Fonds de Recherche du Québec (DOI: https://doi.org/10.69777/309365). The authors thank William Devred (Université Paris-Est Créteil, France) and Chloé Rosa-Teijeiro (Faculté de Médecine Vétérinaire, Université de Montréal) for their contributions to this project. We would also like to thank Prof. Vandesompele (Ghent University - Belgium) for his consistently invaluable advice.

## Author’s contributions

HBM: Conceptualization, Experimentation, Formal analysis, Writing – original draft, Writing – review & editing. FR: Experimentation, Formal analysis, Writing – review & editing. NP: Conceptualization and Experimentation. MHD: Conceptualization, Writing – review & editing, Funding acquisition. FM and MOBB: Project administration, Conceptualization, Validation, Formal analysis, Writing – review & editing, Funding acquisition, Supervision.

## Figure captions

**Figure S1:** Melting curves of the six-candidate reference genes in *Acheta domesticus*. *AdoNEOPT*: AdoNEOPT protein kinase mRNA; *EF1α*: Elongation factor 1-alpha; *EF2*: Elongation factor 2; *GAPDH*: Glyceraldehyde-3-phosphate dehydrogenase; *His H3*: Histone H3; *18S rRNA*: 18S ribosomal RNA.

