## Supplementary figures and images for "Reference Gene Selection for Accurate RT-qPCR Normalization in Four Tissues and Whole-Body Samples of *Acheta domesticus*"

### Melt curves

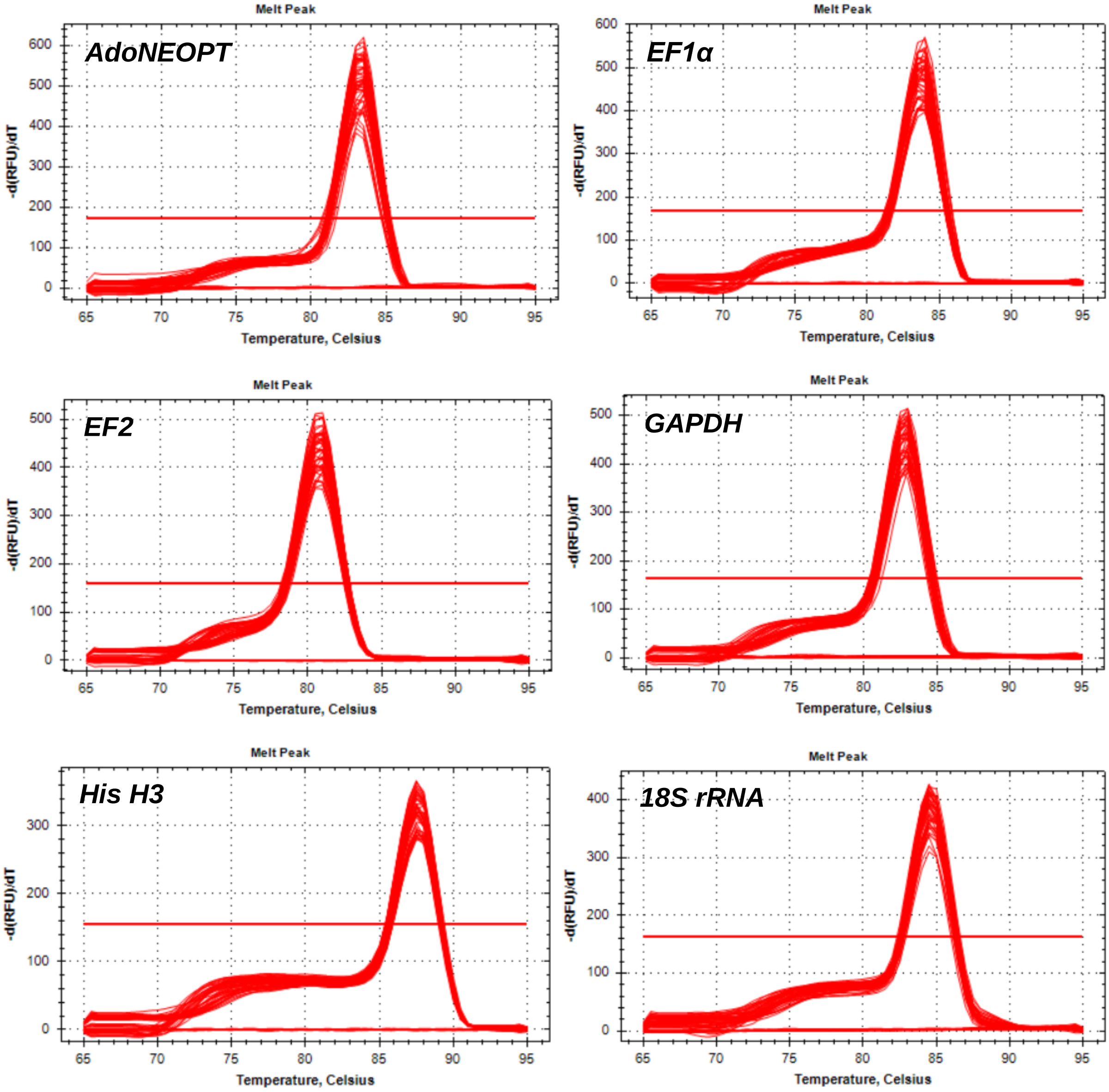
